# Early life food insecurity impairs memory function during adulthood

**DOI:** 10.1101/2025.06.22.660913

**Authors:** Alicia E. Kao, Olivia P. Moody, Emily E. Noble, Kevin P. Myers, Scott E. Kanoski, Anna M. R. Hayes

## Abstract

Approximately 14% of U.S. households are estimated to be food insecure. The neurocognitive and metabolic impacts of unpredictable food access during early-life periods of development are poorly understood. To address these gaps we devised a novel rat model of food insecurity to control the timing, type, and quantity of accessible food using programmable feeders. Male rats were divided into 3 groups: Secure-chow (SC), a control group given 100% of daily caloric needs, distributed evenly across 4 daily meals of standard chow at set mealtimes; Secure-mixed (SM), a 2nd control group identical to the SC group except that the food type predictably alternated daily between chow and a high-fat, high-sugar diet (HFHS); and Insecure-mixed (IM), the experimental group given randomly alternating daily access to either chow or HFHS at either 85% or 115% of daily caloric needs, distributed evenly across 3 daily meals with unpredictable mealtimes. These feeding schedules were implemented from postnatal days (PNs) 26-45, after which all groups received chow ad libitum. Metabolic assessments performed in adulthood revealed no group differences in caloric intake, body weight, or body composition when maintained on either chow (PN46-149) or a cafeteria diet (PN150-174). Behavioral measures (PN66-126) revealed no group differences in anxiety-like, exploratory, or impulsive behavior (zero maze, open field, differential reinforcement of low rates of responding procedures). However, the IM group exhibited hippocampus-dependent memory impairments compared to both control groups in the novel location recognition test. These findings suggest that early-life food insecurity may contribute to long-term impairments in memory function.

## 1. Introduction

Food insecurity (FI) is a state of unpredictable and unreliable access to nutritious food. Experiencing FI during early life is correlated with increased risk of obesity and metabolic disorders (Casey et al., 2006; Dubois et al., 2023; Ortiz-Marrón et al., 2022; Sharkey et al., 2012; Widome et al., 2009; Zhong et al., 2022). In addition to negative metabolic outcomes, there are also correlations in children exposed to FI with cognitive impairments, including deficits in vocabulary, reading, math skills, and emotional problems (Alaimo et al., 2001; de Oliveira et al., 2020; Gallegos et al., 2021; Hobbs & King, 2018; Shankar et al., 2017). In adolescents, FI is associated with greater impulsivity and a higher likelihood of having a mental health disorder such as anxiety, depression, and binge-eating disorder (Reck et al., 2024; Melchior et al., 2012; Ghadban et al., 2025; Bidopia et al., 2023; Nagata et al., 2023). Given the accumulating evidence that FI during development is an important risk factor for adverse metabolic, cognitive, and behavioral outcomes (Dubois et al., 2023; Cohn-Schwartz & Weinstein, 2020; Jyoti et al., 2005), it is critical to develop rigorous experimental models to mechanistically understand whether causal relationships exist between early life FI and these behavioral and physiological outcomes.

Economic, social, psychological, and parental factors modulate the severity and length of early life FI, making it challenging for population studies to isolate specific cause-effect relationships (Berge et al., 2020; Bhargava et al., 2008; Dana et al., 2025; Myers & Temple, 2024; Rahi et al., 2025). Mechanistic research modeling FI in rodents is thus far limited, and results vary significantly regarding the sex, species, and developmental stage studied, as well as the diet type and feeding protocol used (Estacio et al., 2021; Gil et al., 2025; Lin et al., 2022; Myers et al., 2022; Spaulding et al., 2024). Moreover, a defining factor of FI in humans is unpredictability with regards to food accessibility, which includes the amount of food available to consume, the timing of a meal, and the type of food that is available. The majority of these previous rodent models of FI do not utilize unpredictability with regards to individual meal timing, and to our knowledge none of these previous models incorporate food type unpredictability as a variable.

To address these limitations in previous rodent models of FI, we developed a novel early life FI model that strictly targets three defining factors of unpredictable food access: *when food is available*, *how much food is available*, and *what kind of food is available*. During the juvenile and adolescent stages of development, we employed programmable feeders to deliver discrete meals of standard chow or high-fat, high-sugar (HFHS) diet to individually-housed male rats periodically throughout the day. Our model employs two control groups: a group that receives 100% of its average daily caloric needs (i.e., the group as a whole is getting 100% of its overall average daily caloric needs) distributed over 4 daily meals of standard chow given at predictable times (“Secure-chow”, SC), and another group treated identically except that each day the type of food alternates predictably between chow and a HFHS diet (“Secure-mixed”, SM). Our experimental group, which models food insecurity, receives either 85% or 115% of its average daily caloric needs (alternating unpredictably by day) of either chow or HFHS diet (alternating unpredictably by day), and the daily rations are distributed across 3 meals per day that are delivered at unpredictable times (“Insecure-mixed”, IM). Although the IM group’s food quantity that is available to consume purposefully varies by day in this model, total caloric consumption is equated over each 4-day block of feeding to match what both control groups consume, thus making this a model of FI absent changes in total energy consumption.

It is well established that imbalanced nutrition during the juvenile and adolescent stages produces lasting impacts on physical and neurodevelopmental health (Norris et al., 2022; Saavedra & Prentice, 2023). Mechanistic rodent models reveal that different forms of dietary perturbations during these developmental stages, such as malnutrition, undernutrition, or excessive exposure to HFHS “Western” diets, affect brain maturation to produce lasting cognitive and behavioral deficits (Alamy & Bengelloun, 2012; Hayes, Lauer, et al., 2024; Noble & Kanoski, 2016; Tsan et al., 2021). One brain region that is particularly vulnerable to disruption from the early life dietary environment is the hippocampus, which is historically known for its critical role in learning and memory and more recently for its role in appetite and food intake control (Décarie-Spain et al., 2024; Hsu et al., 2018; Kanoski & Grill, 2017; Parent et al., 2022; Rea et al., 2025; Suarez et al., 2020; Yang et al., 2025). We recently identified the period from postnatal (PN) days 26-41 (“early adolescence”) as a particularly sensitive period for diet-induced impacts on hippocampal development, with Western diet consumption limited to the developmental window yielding impairments in hippocampal-dependent learning and memory function that persisted well into adulthood (Hayes, Kao, et al., 2024). Here we seek to expand these previous results to determine whether altered food availability during this critical early adolescence developmental period impacts metabolic outcomes, feeding behavior, behavioral regulation (anxiety-like behavior, impulsivity), and hippocampus-dependent memory using a novel, ethologically-valid, and translational rodent model of FI.

## 2. Methods

### 2.1 Subjects

Male Sprague Dawley rats (N = 46; n = 15-16 per treatment group) from Envigo (Indianapolis, IN, USA) arrived at the University of Southern California animal housing facility on PN day 25 with an average overall initial body weight of 76 g. Animals were individually housed and provided with ad libitum standard chow (Lab Diet 5001; PMI Nutrition International, Brentwood, MO, USA; 29.8% kcal from protein, 13.4% kcal from fat, 56.7% kcal from carbohydrate) and water overnight. Subjects were weighed at 8:30am the following morning (PN 26) to be pseudo-randomly assigned to treatment groups with similar average body weights and were subsequently weighed every morning during scheduled feeding and three times a week during ad libitum feeding. All subjects were individually housed from arrival through PN 98 when they were subsequently double-housed with a member from their own treatment group (one subject per treatment group remained single-housed from PN 98 onward due to uneven subject numbers per group). The housing room was climate-controlled and on a 12:12 h reverse light/dark cycle with lights turned off at 11am and turned on at 11pm. All experimental procedures were approved by the University of Southern California Institutional Animal Care and Use Committee (IACUC) following the National Research Council Guide for the Care and Use of Laboratory Animals.

### 2.2 Scheduled feeding protocols

Juvenile male rats were maintained on the scheduled feeding protocol for 20 days from PN 26-45 before being transferred to ad libitum standard chow (Lab Diet 5001; Fig. 1A). The ad libitum chow given upon arrival to the animal housing facility on PN 25 was removed at 8:30am on PN 26. Animals received daily rations of food that were distributed three or four times per day (depending on treatment group) at 11am, 3pm, 7pm, and/or 11pm by programmable automated 3D printed feeders. Each home cage was equipped with its own feeder that dropped meals from a rotating carousel directly onto a wire rack in the cage that was readily accessible to the rats. The feeders were programmed using an Arduino Nano microcontroller with a real-time clock chip which actuates a 12V stepper motor to rotate the carousel. To predetermine daily rations of food for the growing animals, we matched the average weight of all animals on PN 26 with archival data from ∼70 previous juvenile male Sprague Dawley rats in our lab. The data was averaged across this developmental period to generate the average daily caloric targets.

**Figure 1.**
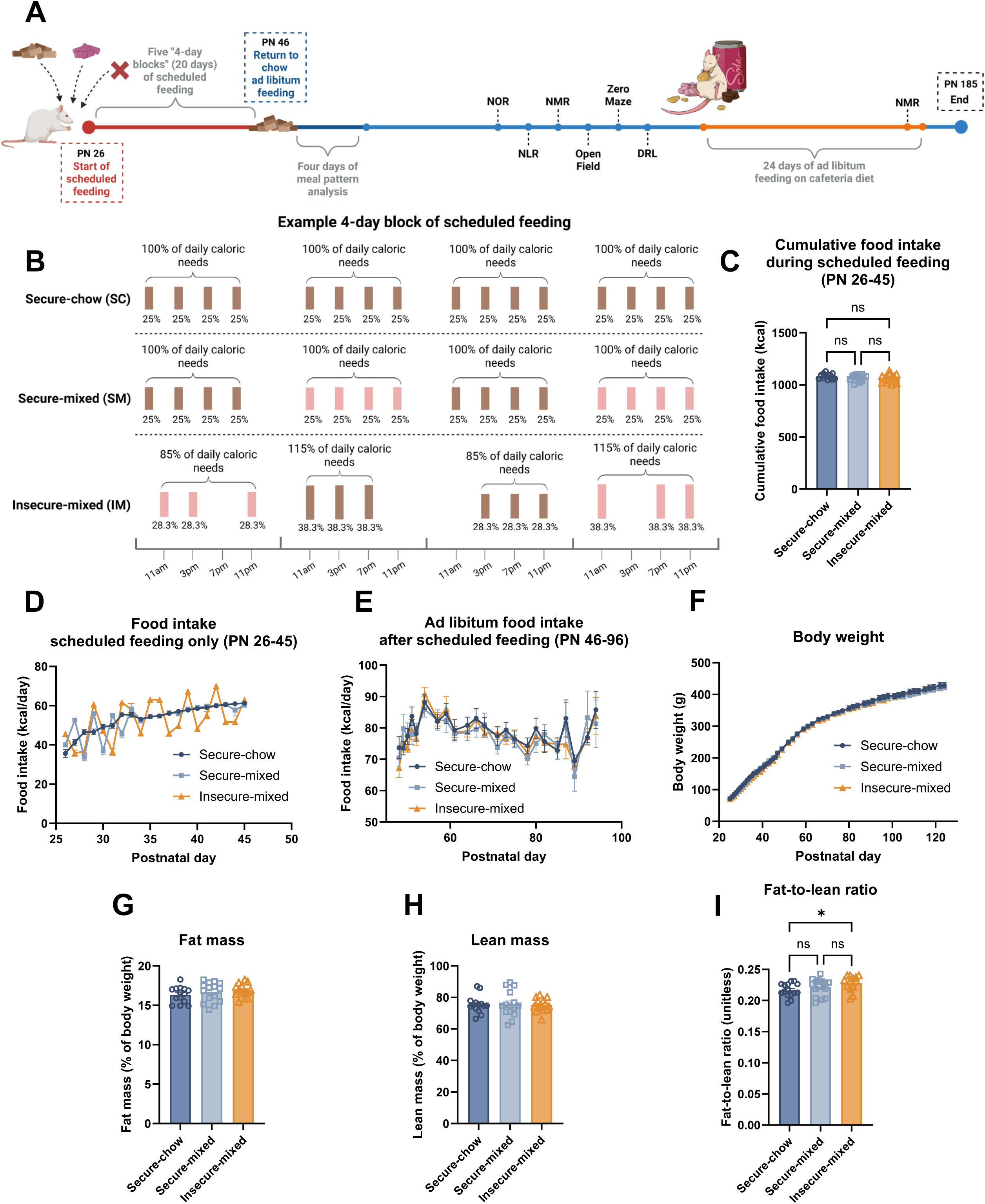
Male rats exposed to early life food insecurity do not exhibit differences in food intake, body weight, or body composition in adulthood. A: Timeline of experimental procedures. B: Example of one 4-day block of the scheduled feeding period (PN 26-45) for each of the three groups, illustrating how total all groups received 100% of their caloric needs per block. C: Cumulative food intake during the scheduled feeding period (one-way ANOVA [group]; P=0.27 overall). D: Daily food intake over time during the scheduled feeding period (mixed effects model with repeated measures [group, time, group × time interaction]; P=0.51 for group, P<0.0001 for time, P<0.0001 for group × time interaction; many post hoc differences using Tukey’s multiple comparisons test due to the nature of the food insecurity model). E: Ad libitum food intake over time after the scheduled feeding period (mixed effects model with repeated measures [group, time, group × time interaction]; P=0.78 for group, P<0.0001 for time, P=0.99 for group × time interaction). F: Body weight over time (mixed effects model with repeated measures [group, time, group × time interaction]; P=0.85 for group, P<0.0001 for time, P<0.99 for group × time interaction). G: Fat mass as a percentage of body composition in adult rats after early life food insecurity (one-way ANOVA [group]; P=0.38). H: Lean mass as a percentage of body composition in adult rats (one-way ANOVA [group]; P=0.97). I: Fat-to-lean ratio of body composition in adult rats (one-way ANOVA [group]; P=0.0587, c). n=14-16 per group. *P <0.05. DRL, differential reinforcement of low rats of responding procedure; NLR, novel location recognition task; NOR, novel object recognition task; NMR, nuclear magnetic resonance measure of body composition; PN, postnatal day.

The FI model had three treatment groups, matched in overall amount of accessible food distributed in each of five 4-day “blocks” and thus also matched for the total amount fed during the 20-day feeding protocol. There were two control groups: the Secure-chow (SC) group received standard chow (Rodent Diet 5001) every day, while the Secure-mixed (SM) group received the same amount of food (in kcal), alternating daily between standard chow and high fat, high sugar diet (HFHS diet; Research Diets D12415, Research Diets Inc., New Brunswick, NJ, USA; 20% kcal from protein, 35% kcal from carbohydrate, 45% kcal from fat). Both groups received 100% of their predetermined caloric needs each day based on archival data, with food rations evenly divided across four meals delivered at the set mealtimes previously described.

To model FI, within each 4-day block of the scheduled feeding period, the Insecure- mixed (IM) group had two pseudo-randomly selected days when they received 85% of the block’s average daily food ration (low food days) and two pseudo-randomly selected days when they received 115% of the block’s average daily food ration (high food days). This schedule ensured that at the end of the four days, average caloric intake equated to be 100% of what the SC and SM groups consumed within that same block (i.e., overall caloric intake was matched across groups within each block). Independent of food quantity assignments, diet type was also manipulated so that two days out of each 4-day block were pseudo-randomly selected to be standard chow or HFHS days, with equivalent caloric content of the rations between diet types (due to differing energy densities between standard chow and HFHS). The schedule of high and low food days, along with chow and HFHS days, was designed so that there were no more than two consecutive days of either quantity or diet type. The IM group’s daily rations were also evenly divided into three meals with one of the four mealtimes skipped each day to integrate the variable of unpredictable meal timing. Each mealtime was randomly skipped once during each 4-day block. An example of a single 4-day feeding block is presented in Fig. 1B.

Over the course of the scheduled feeding period, cages, bedding, and wire racks were carefully monitored for any leftover portions of food. If any food was left before the start of the next day’s feeding (i.e., before the 11am scheduled feeding), it was removed and weighed for accurate recording of food intake. All subjects had free access to water throughout the duration of the scheduled feeding period.

### 2.3 Ad libitum feeding

#### 2.3.1 Ad libitum chow intake

Immediately following the 20-day scheduled feeding period, on PN 46, animals were transferred to a BioDAQ food intake monitoring system (Research Diets Inc., New Brunswick, NJ), with ad libitum standard chow and water. Food intake parameters including daily food intake, number of meals, meal size, and meal duration were continuously recorded. Meals were defined as food consumption greater than 0.1g with eating bouts within 10 minutes of each other considered part of the same meal. Meal pattern analyses were performed for data collected from PN 49-52, allowing for an initial habituation period to the food intake monitoring system.

On PN 53-54, animals were transferred back to standard home cages and were maintained on ad libitum standard chow and water for all behavioral testing (Figure 1A). Chow intake was measured three times a week.

#### 2.3.2 Ad libitum consumption of a “junk food” cafeteria diet challenge in adulthood

From PN 152-174, a total of 29 rats (n = 10 SC group, n = 10 SM group, n = 9 IM group) were transferred to hanging wire cages and challenged with an ad libitum “junk food” style cafeteria diet (CAF) as detailed previously (Hayes, Lauer, et al., 2024; Hayes et al., 2022). The CAF diet was composed of HFHS diet, potato chips (Ruffles Original, Frito Lay, Casa Grande, AZ, USA), chocolate-covered peanut butter cups (Reese’s Minis Unwrapped, The Hershey Company, Hershey, PA, USA), and 11% weight/volume high-fructose corn syrup-55 (HFCS) beverage (Best Flavors, Orange, CA, USA) alongside water. Each CAF diet component was placed in separate food receptacles or sipper bottles and individually weighed three times a week shortly before the onset of the dark cycle (9:00-10:30am). Cardboard sheets were placed underneath each cage to accumulate spillage which was also collected and measured during weighing days to allow for accurate assessment of food intake. Energy (kcal) consumed from each of the CAF diet components was calculated by multiplying the measured weight of food/drink consumed per rat by the energy density of the respective component (4.7 kcal/g for HFHS diet, 5.7 kcal/g for potato chips, 5.1 kcal/g for peanut butter cups, 0.0296 kcal/g for HFCS beverage).

### 2.4 Nuclear magnetic resonance (NMR) assessment of body composition

Body composition parameters (percent fat mass, percent lean mass, fat-to-lean ratio) were assessed using a Bruker NMR Mini-spec LF 90II (Bruker Daltonics, Inc., Billerica, MA, USA) at PN 82 (following the scheduled feeding period and some behavior assessments) and for a subgroup of rats at PN 173 (during the CAF diet period). Rats were food restricted for 1 h and weighed prior to scanning. Percent fat mass and percent lean mass were calculated as [fat mass (g)/body weight (g)] × 100 and [lean mass (g)/body weight (g)] × 100, respectively.

### 2.5 Novel object recognition (NOR)

NOR is a behavioral assay that evaluates preferential exploration of a novel object to assess perirhinal cortex-dependent recognition memory (Albasser et al., 2011). NOR was tested in a dimly lit room using a grey opaque plastic box (38.1 cm L × 56.5 cm W × 31.8 cm H) flanked by desk lamps on two opposing sides of the box. Light was directed towards the floor to produce soft, ambient lighting. Rats were placed into the empty box for 10 min to habituate to the apparatus 1-2 days prior to testing. On test day, rats were placed in the center of the box, now containing two identical objects. Rats were initially positioned to face a neutral wall in order to avoid biasing them toward either object. The objects used were either two identical stemless wine glasses or two identical ceramic brown jugs. The rats were allowed to freely explore for 5 min to familiarize themselves with the objects (‘familiar objects’) before being removed and placed in their home cage for 5 min. During this period, the box and objects were cleaned with 10% ethanol solution, and one of the duplicate objects was replaced with a novel object (whichever object the animal had not been previously exposed to). Rats were then placed back into the center of the box and allowed to freely explore for 3 min. The side where the novel objects were placed, as well as the assigned novel and familiar objects themselves were counterbalanced by treatment group. An experimenter blinded to group and novel object assignments used video recordings to hand-score the total time that each rat spent actively exploring each object, defined as sniffing or touching the object with the nose or forepaws. Exploration index was calculated as [time spent exploring novel object / total time spent exploring both objects] during the 3-min test exposure.

### 2.6 Novel location recognition (NLR)

NLR is a test of hippocampal-dependent spatial working memory (Denninger et al., 2018) using the same box apparatus, habituation protocol, and lighting conditions as NOR. On test day, rats were placed in the center of the box containing two identical objects placed in a horizontal line along the length of one wall. The objects used were either two identical soap dispensers or two identical saltshakers (all objects were empty and clean). The rats were allowed to freely explore for 5 min to familiarize themselves with the objects before being removed and placed in their home cage for 5 min. During this time, the box and the objects were cleaned with 10% ethanol solution. One of the objects was moved to a different location (‘object in a novel location’), forming a diagonal line across the box. Rats were placed back into the center of the box and allowed to freely explore for 3 min. The wall in which the objects were initially lined up against, the novel location, and the set of objects used were counterbalanced by treatment group. An experimenter blinded to group and novel location assignments used video recordings to hand-score the total time that each rat spent actively exploring each object. Exploration index was calculated as [time spent exploring object in a novel location / total time spent exploring both objects] during the 3-min test exposure.

### 2.7 Zero maze

The elevated zero maze is a circular runway designed to evaluate anxiety-like behavior (Belzung & Griebel, 2001). The ring-shaped track (11.4 cm wide track, 73.7 cm height from track to the ground, 92.7 cm exterior diameter) was divided into four equal length quadrants that alternate between two open segments with 3-cm high curbs and two closed segments with 17.5 cm high walls. Rats were placed in the middle of an open segment positioned parallel with the track and allowed to freely roam for 5 min. After each test session, the apparatus was cleaned with 10% ethanol. ANYmaze activity tracking software (Stoelting Co., Wood Dale, IL, USA) was used with video recordings to measure the total amount of time each animal spent in the open or closed segments as well as the number of entries made into the respective segments.

### 2.8 Open field

The open field test measures locomotor activity and evaluates anxiety-like behavior in an open arena (Belzung & Griebel, 2001). The apparatus consisted of a grey opaque plastic box (53.5 cm L × 54.6 cm W × 36.8 H) with a rectangular center zone (19 cm × 17.5 cm). Five desk lamps were placed around the apparatus to produce diffuse brighter lighting (∼44 lux) in the center zone and darker lighting (∼30 lux) in the corners of the box. Rats were placed in the center of the box and allowed to freely roam for 10 min. After each test session, the apparatus was cleaned with 10% ethanol. ANYmaze activity tracking software (Stoelting Co., Wood Dale, IL, USA) was used with video recordings to measure total distance travelled within the apparatus (locomotor activity) along with total time spent in the illuminated center zone (measure of anxiety-like behavior).

### 2.9 Differential reinforcement of low rates of responding (DRL)

DRL is an operant procedure designed to assess impulsive action towards a reward (Noble et al., 2019). Impulsive responses are evaluated by the success rate of subjects to withhold lever presses for pre-determined amounts of time in exchange for receiving a high-fat palatable reward (45 mg dustless precision pellets; 35% kcal fat and sucrose-enriched, F05989, Bio-Serv, Flemington, NJ, USA). Rats were first habituated in their home cage to the reward pellets before starting DRL. On each day of DRL training, chow was removed from the home cage 1 h before training sessions, which started at the onset of the dark cycle and lasted 45 min. Rats were trained in operant chambers (Med Associates, Fairfax, VT, USA) containing an active lever (reinforced press dispensed one pellet reward with lever light activation) and an inactive lever (non-reinforced). For the first 3+ days of training, animals were on a DRL0 schedule in which active lever presses were immediately reinforced with a 0-s time delay and no reward limit. Animals were then switched to a DRL5 schedule for 3+ days, during which they were trained to withhold lever presses for at least 5 s after each press before subsequent lever presses could be rewarded. If either the active or inactive lever was pressed within the withholding period, the timer restarted. DRL 5 was followed immediately by 4 days of a DRL10 schedule (10 s withholding period) and 10 days of a DRL20 schedule (20 s withholding period). Efficiency was calculated as [number of reinforced lever presses / number of active lever presses].

### 2.10 Statistical analyses

Statistical analyses and figure generation were performed using Prism software (GraphPad, Inc., version 10.4.1, San Diego, CA, USA). Data are presented as mean ± standard error of the mean (SEM) for error bars in all figures, and significance was considered at P < 0.05. Specific statistical tests and analyses results for each figure panel are detailed in Supplementary Table S1 along with the number of subjects used for each experiment.

## 3. Results

### 3.1. Early life food insecurity does not alter subsequent ad libitum food intake, body weight, or body composition

The early life food insecurity model effectively altered the pattern of daily food intake in the IM group while, as expected, not altering overall cumulative food intake between groups during the scheduled feeding period (P=0.27 for cumulative food intake during the scheduled feeding period, Fig. 1C; P<0.0001 for interaction of group × time with significant post-hoc differences at all but two time points, Fig. 1D). Food intake from PN 26-33 during the scheduled feeding period (i.e., during the first two blocks) was more variable between groups because our estimates for projected food intakes from archival data were initially ∼18% too high. This was evident because rats in both the SM and IM groups ate a majority of the rations on HFHS days but left some of the rations on chow days (Fig. 1D). We adjusted our estimates for projected food intakes for blocks 2-5 (and therefore reduced the food rations given, matching the amounts within blocks) and the feeding schedule was implemented as intended from PN33-45 (Fig. 1D). No differences in ad libitum chow intake were observed between groups after the scheduled feeding period (P=0.78 for main effect of group, P<0.0001 for main effect of time [indicating growth], and P=0.99 for the interaction of time × group; Fig. 1E). Body weight did not significantly differ between groups during the scheduled feeding period or subsequent ad libitum chow intake period (P=0.85 for main effect of group, P<0.0001 for main effect of time [indicating growth], and P>0.99 for the interaction of time × group; Fig. 1F). Although there were no differences in body composition outcomes of fat mass or lean mass between groups (P=0.38 for fat mass, P=0.97 for lean mass; Fig. 1G-H), there was a trending overall difference in fat-to-lean ratio accompanied by a significant post hoc comparison between the SC and IM groups (P=0.059 for main effect, P=0.047 for post hoc Tukey’s multiple comparisons test between SC and IM; Fig. 1I).

### 3.2. Food insecurity in early life does not modify meal patterns of subsequent ad libitum chow consumption

Despite the lack of differences in overall food intake between groups upon the switch to ad libitum chow consumption following the scheduled feeding period, we hypothesized that early life FI may still have shown altered meal structure. To test this hypothesis, on PN 46 (the day after the scheduled feeding period ended) the rats were placed into a comprehensive food intake monitoring system for the measurement of meal patterns from PN 49-52. Contrary to our hypothesis, no differences were observed for daily food intake, number of meals, meal duration, or meal size (P=0.72, P=0.18, P=0.16, P=0.61, respectively; Fig. 2A-D). In addition, no differences were found for size, eating rate, or duration of the first meal per day during the measurement period (P=0.61, P=0.99, P=0.94, respectively; Fig. 2E-G).

**Figure 2.**
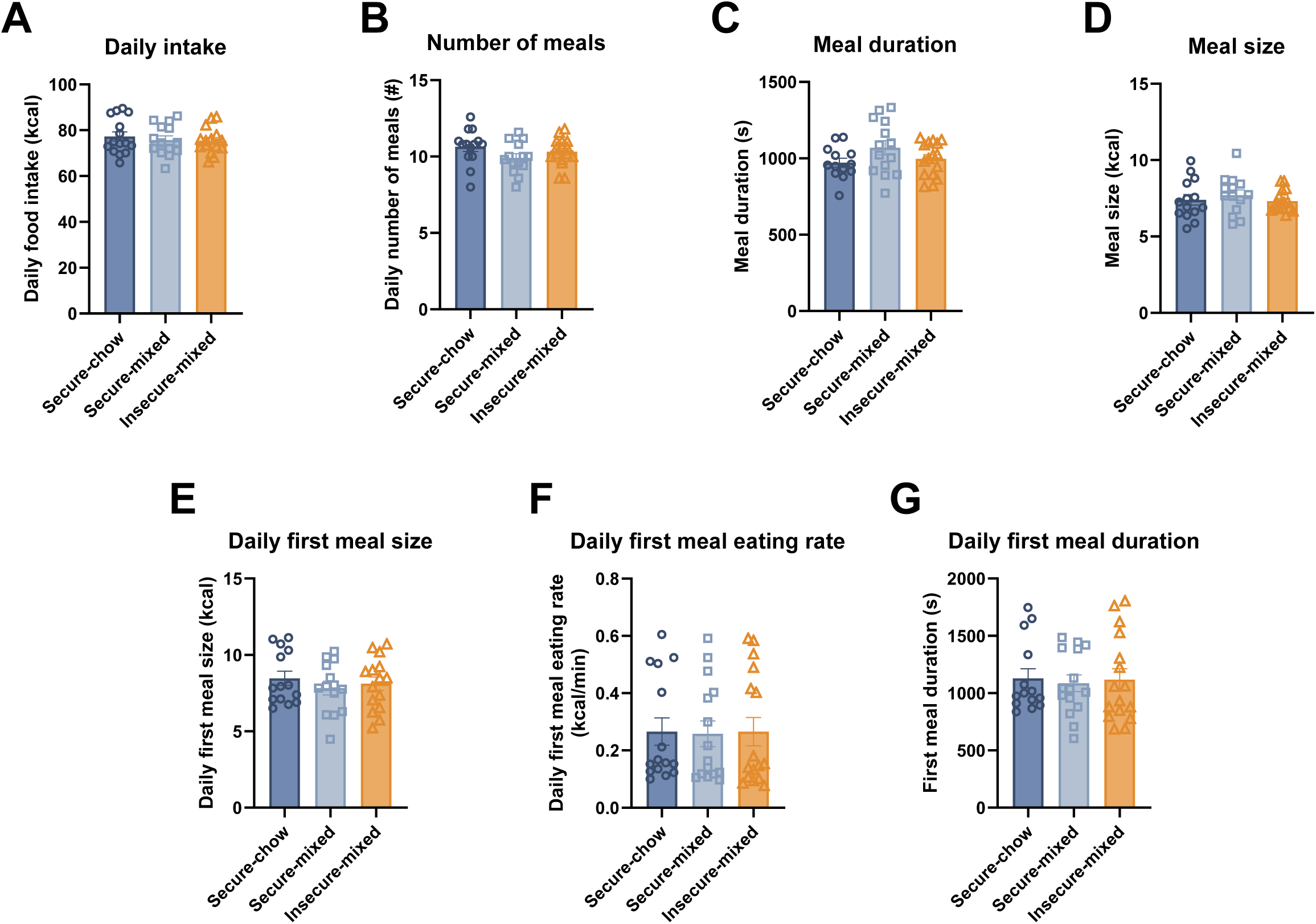
Early life food insecurity in male rats does not alter meal patterns in the short-term when subsequently switched to ad libitum standard chow consumption. A: Daily food intake of standard chow measured by an automatic food intake monitoring system shortly after the end of the scheduled feeding period (one-way ANOVA [group]; P=0.72). B: Number of meals consumed per day shortly after the end of the scheduled feeding period (one-way ANOVA [group]; P=0.18). C: Meal duration shortly after the end of the scheduled feeding period (Brown-Forsythe ANOVA [group] due to unequal standard deviation between groups; P=0.16). D: Meal size shortly after the end of the scheduled feeding period (one-way ANOVA [group]; P=0.61). E: Daily first meal size shortly after the end of the scheduled feeding period (one-way ANOVA [group]; P=0.61). F: Daily first meal eating rate shortly after the end of the scheduled feeding period (one-way ANOVA [group]; P=0.99). G: Daily first meal duration shortly after the end of the scheduled feeding period (one-way ANOVA [group]; P=0.94). n=14-16 per group.

### 3.3. Early life food insecurity impairs hippocampus-dependent memory function but not perirhinal cortex-dependent object novelty memory during adulthood

In the novel location recognition (NLR) test, rats in the IM group did not preferentially explore an object moved to a novel location relative to an object that remained in a familiar location (P=0.02 for exploration index during the test phase [Tukey’s post hoc multiple comparisons: P=0.98 for SC vs. SM, P=0.04 for SC vs. IM, and P=0.04 for SM vs. IM], P=0.95 for total object exploration time during initial exposure; Fig. 3A-C), signifying an impairment in hippocampus-dependent memory for the IM group relative to the SC and SM groups. Further, both the SC and SM groups had exploration indices that differed significantly from chance during the test phase, whereas the IM group’s exploration index did not differ from chance (P<0.0001 for SC and SM groups vs. P=0.44 for IM group, one-sample t-test as a comparison of each group’s actual mean to the theoretical mean of 0.5 [indicative of chance exploration of both objects]), which indicates that the SC and SM groups remembered previous object locations whereas the IM group did not.

**Figure 3.**
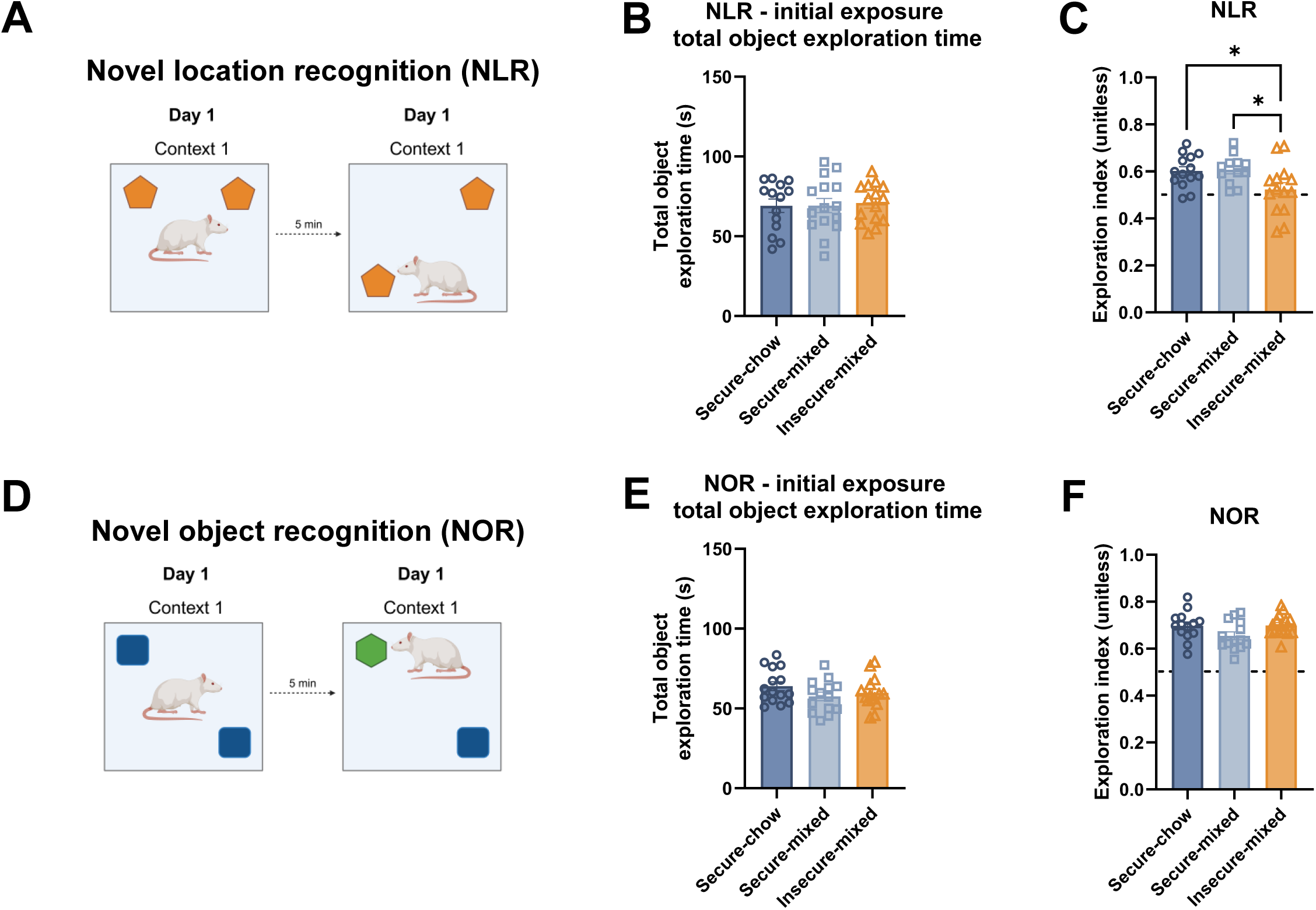
Early life food insecurity impairs hippocampus-dependent spatial recognition memory, but not object novelty recognition during adulthood in male rats. A: Diagram of NLR task. B: Total object exploration time during the initial exposure stage of NLR (one-way ANOVA [group]; P=0.95). C: NLR object exploration index during the test stage (one-way ANOVA [group]; P=0.02, post hoc Tukey’s multiple comparisons tests for SC vs. SM, P=0.98; SC vs. IM, P=0.04; SM vs. IM, P=0.04). D: Diagram of NOR task. E: Total object exploration time during the initial exposure stage of NOR (one-way ANOVA [group]; P=0.23). E: NOR object exploration index during the test stage (one-way ANOVA [group]; P=0.07). n=12- 16 per group. *P <0.05. NLR, novel location recognition; NOR, novel object recognition.

No differences were found in the novel object recognition NOR test, which relies on the perirhinal cortex to facilitate object-novelty recognition (P=0.23 for total object exploration time during initial exposure, P=0.07 for exploration index during the test phase; Fig. 3D-F). All groups exhibited learning in the NOR task (P<0.0001 for exploration index for the SC, SM, and IM groups; one-sample t-test as a comparison of each group’s actual mean to the theoretical mean of 0.5 [indicative of chance exploration of both objects]). These results suggest that the impairment in NLR in the IM group was not based on novelty avoidance.

### 3.4. Exposure to food insecurity in early life does not alter anxiety-like behavior, locomotor activity, or impulsivity during adulthood

In the zero maze test for anxiety-like behavior, no differences were observed between groups for time spent in the open arms or number of entries into the open arms (P=0.30 and P=0.81, respectively; Fig. 4A-C). In the open field test for anxiety-like behavior and locomotor activity, no differences were found between groups for the amount of time spent in the center of the arena (P=0.09; Fig. 4D-E). A significant effect of group was found for the amount of distance travelled during the open field test (P=0.01; 4F), and post hoc tests indicated that the SM group travelled a significantly greater distance than the SC group (Tukey’s post hoc multiple comparisons: P=0.01 for SC vs. SM). However, there were no differences in the amount of distance travelled between the IM and either SC or SM control groups (Tukey’s post hoc multiple comparisons: P=0.34 for SC vs. IM, P=0.25 for SM vs. IM). No differences were found in impulsive behavior between groups, as evidenced by lack of differences in DRL 20 efficiency (P=0. 87 for main effect of group, P<0.0001 for main effect of time [indicating learning], and P=0.63 for the interaction of time × group; Fig. 4H).

**Figure 4.**
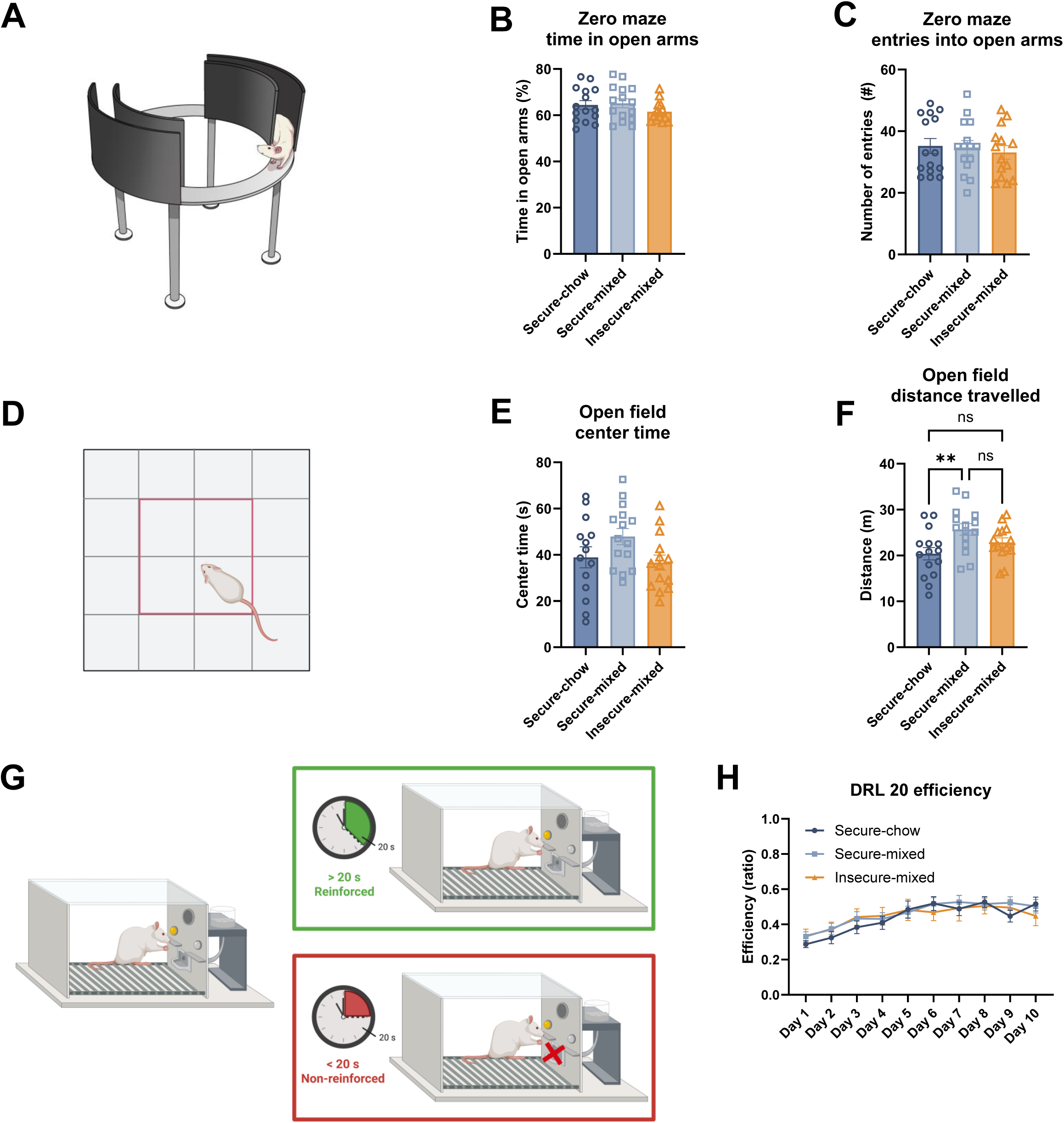
Exposure to food insecurity in early life does not alter anxiety-like behavior, locomotor activity, or impulsivity in male rats during adulthood. A: Zero maze apparatus diagram. B: Time spent in the open arms during the zero maze task (one-way ANOVA [group]; P=0.30). C: Entries into the open arms during the zero maze task (one-way ANOVA [group]; P=0.81). D: Diagram of the open field task. E: Time spent in the center zone of the open field test (one-way ANOVA [group]; P=0.09). F: Distance travelled during the open field test (one-way ANOVA [group]; P=0.01, post hoc Tukey’s multiple comparisons tests for SC vs. SM, P=0.01; SC vs. IM, P=0.34; SM vs. IM, P=0.25). G: DRL protocol schematic. H: Efficiency of rats in performing (learning) DRL when an interval of 20 s with no lever presses was required in order to earn a reward (food pellet; two-way ANOVA with repeated measures [group, time, group × time interaction]; P=0.87 for group, P<0.0001 for time [indicating learning], P=0.63 for group × time interaction). n=14/15 per group. *P <0.05. DRL, differential reinforcement of low rates of responding procedure.

### 3.5. Early life food insecurity does not influence responses to a cafeteria-style junk food diet challenge in adulthood

Body weights did not significantly differ between groups upon 24-day ad libitum consumption of a high-fat high-sugar CAF diet from PN 150-174 (P=0.36 for main effect of group, P<0.0001 for main effect of time, and P=0.38 for the interaction of time × group; Fig. 5A). Furthermore, there were no differences in total energy intake between groups during the CAF diet period (P=0.74 for main effect of group, P<0.0001 for main effect of time, and P=0.15 for the interaction of time × group; Fig. 5B). While groups did not differ in their consumption of the HFHS chow component of the CAF diet (P=0.79 for main effect of group, P<0.0001 for main effect of time, and P=0.55 for the interaction of time × group; Fig. 5C), there was a fleeting difference in intake of the potato chip component of the CAF diet, with the SM group consuming significantly fewer kcal from potato chips at PN 152 (P=0.28 for main effect of group, P<0.0001 for main effect of time, and P=0.01 for the interaction of time × group; post hoc differences at PN 152 only using Tukey’s multiple comparisons test: P=0.02 for SC vs. SM, P=0.97 for SC vs.

**Figure 5.**
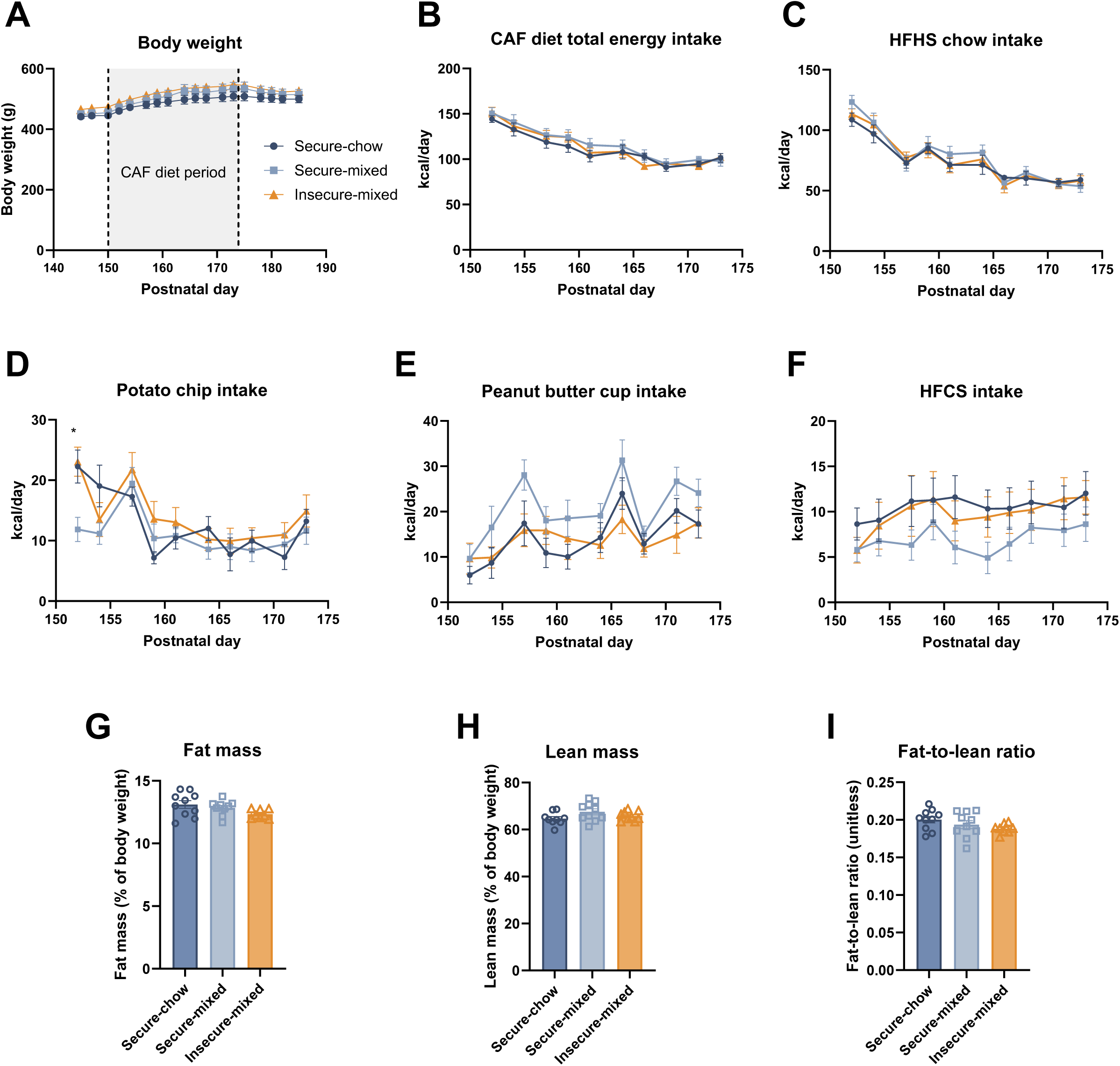
Early life food insecurity did not alter food intake or body composition upon a high-fat, high-sugar junk food cafeteria diet challenge during adulthood. A: Body weight over time during the cafeteria diet challenge period (two-way ANOVA with repeated measures [group, time, group × time interaction]; P=0.36 for group, P<0.0001 for time [indicating weight gain], P=0.38 for group × time interaction). B: Total energy intake during the cafeteria diet period (two-way ANOVA with repeated measures [group, time, group × time interaction]; P=0.74 for group, P<0.0001 for time [indicating weight gain], P=0.15 for group × time interaction). C: Energy intake from the high fat high sugar diet component during the cafeteria diet period (two-way ANOVA with repeated measures [group, time, group × time interaction]; P=0.79 for group, P<0.0001 for time [indicating weight gain], P=0.55 for group × time interaction). D: Energy intake from the potato chip component during the cafeteria diet period (two-way ANOVA with repeated measures [group, time, group × time interaction]; P=0.28 for group, P<0.0001 for time [indicating weight gain], P=0.01 for group × time interaction; post hoc Tukey’s multiple comparisons tests indicated significant differences between groups only at PN 152: SC vs. SM, P=0.01; SC vs. IM, P=0.97; SM vs. IM, P=0.01). E: Energy intake from the peanut butter cup component during the cafeteria diet period (mixed effects model with repeated measures [group, time, group × time interaction]; P=0.04 for group, P<0.0001 for time, P=0.67 for group × time interaction). F: Energy intake from the high- fructose corn syrup component during the cafeteria diet period (two-way ANOVA with repeated measures [group, time, group × time interaction]; P=0.37 for group, P=0.06 for time, P=0.90 for group × time interaction). G: Fat mass as a percentage of body composition in adult rats after 23 days of the cafeteria diet challenge (Brown-Forsythe ANOVA [group] due to unequal standard deviation between groups; P=0.08). H: Lean mass as a percentage of body composition in adult rats after 23 days of the cafeteria diet challenge (one-way ANOVA [group]; P=0.13). I: Fat-to-lean ratio of body composition in adult rats after 23 days of the cafeteria diet challenge (one-way ANOVA [group]; P=0.19, c). n=8-10 per group. *P <0.05. CAF, cafeteria diet; HFCS, high-fructose corn syrup; HFHS, high fat high sugar chow; PN, postnatal day.

IM, P=0.01 for SM vs. IM; Fig. 5D). There was a main effect of group for consumption of peanut butter cups, with the SM group exhibiting greater overall kcal intake from this CAF diet component relative to the SC and IM groups (P=0.04 for main effect of group, P<0.0001 for main effect of time, and P=0.67 for the interaction of time × group; Fig. 5E). No differences in kcal consumption of the HFCS beverage were found between groups (P=0.37 for main effect of group, P=0.06 for main effect of time, and P=0.90 for the interaction of time × group; Fig. 5E). Additionally, there were no differences in fat mass, lean mass, or fat-to-lean ratio measures of body composition between groups after consuming the CAF diet for 23 days (P=0.08, P=0.13, P=0.19, respectively; Fig. 5G-I).

## 4. Discussion

Despite its rising prevalence in the United States (Rabbitt et al., 2024), little is understood about how experiencing food insecurity (FI) during early life impacts juvenile development. Population studies reveal that unstable access to food during childhood is linked with poor cognitive and health outcomes (Fleming et al., 2021; Frongillo et al., 2024; Gallegos et al., 2021; Shankar et al., 2017; St Pierre et al., 2022), yet causal effects on neural and metabolic development have not been well established from these correlational findings. To mechanistically study causal relationships between FI and metabolic and behavioral outcomes using a rodent model, we devised a novel scheduled-feeding protocol using automated programmable feeders that exacts full experimental control over the quantity, timing, and type of food delivered to each subject over 20 critical days of juvenile development. Importantly, monitoring of food consumption during the 20-day early life scheduled feeding period confirmed that groups did not differ in cumulative caloric intake, as intended in the implementation of the model, and thus allowing for inferences to specifically be made about the unpredictable, irregular nature of FI in the absence of food restriction. Once switched to ad libitum feeding, no group differences were found in food intake, body weight, body composition, or feeding behavior. Although juvenile FI did not affect anxiety-like behavior, locomotion, impulsive responses to food reward, or perirhinal cortex-dependent object novelty recognition during adulthood, it did impair hippocampal-dependent spatial memory in the novel location recognition procedure. These collective findings highlight the hippocampus as being especially vulnerable to unpredictable dietary stressors during early development.

Food insecurity can have differing levels of severity. The USDA defines two levels of food insecurity: ‘low food security’ and ‘very low food security’ (USDA Economic Research Service). A notable distinction between these levels is food adequacy, with ‘very low food security’ being marked by reduced food intake (inadequate food) and ‘low food security’ marked by little or no indication of reduced food intake (adequate food). ‘Low’ food security is more prevalent than ‘very low’ food security in the United States, especially among children and adolescents (Rabbitt et al., 2024), therefore our aim was specifically to model the effects of unpredictable access to food in the absence of undernutrition. We employed two control groups that received either standard chow or a mixed diet of standard chow and HFHS diet to identify outcomes specific to unpredictable food access. The Secure-chow (SC) group established the baseline outcomes produced by highly regulated meal deliveries instead of free, unrestricted feeding, and the Secure-mixed (SM) group was used to match the diet composition of the Insecure-mixed (IM) group.

Body weight did not differ between groups throughout the experiment, a finding that is difficult to compare with existing animal studies in the literature, as each study’s FI model is markedly distinct. Few models have purposefully controlled for caloric intake during unpredictable food access, and there has yet to be consistent replication of findings across different developmental stages, the sex studied, and the type of diet used. Notably, prior research with FI models using standard chow indicates that food insecure mice consume fewer calories during FI yet have similar body weights to control groups (Estacio et al., 2021; Gil et al., 2025). In FI models that incorporated a high-fat diet, Spaulding and colleagues (2024) found no food intake or body weight differences during or after a juvenile FI period, while Myers and colleagues (2022) reported rapid weight gain in food insecure adult female rats once provided with ad libitum access. Collectively, these observations suggest that in rodent models of FI, differential weight gain depends on an interaction between food unpredictability and some other factors of the protocol, which may include diet type, age, and/or sex.

Similar to how existing models of FI widely vary, children’s experiences of FI are also inherently diverse. This variability has produced inconsistent findings regarding FI’s impact on weight status and obesity risk. FI among children has been linked with both increased (Eskandari et al., 2022; Fleming et al., 2021; Ortiz-Marrón et al., 2022) and decreased (Matheson et al., 2002; Rose & Bodor, 2006; Wirth et al., 2020) odds of childhood overweight or obesity status, and numerous review articles highlight a similar mix of positive and negative findings (Eisenmann et al., 2011; Frongillo et al., 2024). Along with these varying associations between childhood FI and current weight status, longitudinal studies suggest that early life FI leads to increased weight gain over time, even throughout young adulthood (Dubois et al., 2023; Liu et al., 2025; Metallinos-Katsaras et al., 2012; Zhong et al., 2022). However, evidence suggests that there are likely to be important sex differences implicated in the impacts of early life FI on body weight, as food-insecure girls face higher obesity risk compared to boys (St Pierre et al., 2022). Support for such sex differences has been borne out in rodents, as food-insecure juvenile female mice exhibit increased body weight gain, whereas juvenile males do not (Lin et al., 2022). While the present study was limited by only using males, revealing no impact of FI on energy intake and body weight (consistent with previous literature), our future follow-up research will evaluate the impact of this novel rodent model of FI on metabolic and cognitive outcomes in female rats.

Previous rodent studies have found that FI leads to increases in fat mass (Gil et al., 2025; Spaulding et al., 2024). FI has also been studied in other model organisms such as the European Starling, showing greater weight gain and fat stores in response to FI due to increased energetic efficiency and not from increased caloric intake (Andrews et al., 2021; Bateson et al., 2021).

Similarly, early life FI in humans is associated with increased fat mass and risk of obesity throughout the lifespan (Dubois et al., 2023; Liu et al., 2025; Metallinos-Katsaras et al., 2012; Zhong et al., 2022). Nettle and colleagues (2017) outline an “insurance hypothesis,” proposing an evolutionary perspective that FI adaptively favors increased fat storage. During periods of unpredictable food access, weight gain in the absence of increased energy intake likely serves as a protective mechanism against future food deprivation and supports a link between FI and obesity. However, in our model we did not observe differences in food intake, body weight, or fat percentage when rats were maintained ad libitum either on chow or an energy-dense cafeteria diet, suggesting that the effects of FI on fat mass may be sex- and/or species-dependent.

Another adaptive strategy that may be used in response to FI is overconsumption during abundant food access and developing a preference for calorically dense foods. Rodents acutely increase food intake following food deprivation (Tagliaferro & Levitsky, 1982), which was observed during recovery periods following restriction in most FI models, including the present model. However, it is unclear whether a history of FI, during development or adulthood, encourages persistent increased food intake once stable food access is restored. Spaulding and colleagues (2024) and our current study collectively demonstrate in rats that exposure to FI during development does not necessarily lead to higher food intake of either standard chow or an array of high fat, high sugar foods in adulthood.

There is also evidence that adolescents experiencing FI have altered eating behavior and food choices. Food-insecure children are found to have increased fat intake, eat more fast food, and have lower vegetable consumption (Grutzmacher & Gross, 2011; Sharkey et al., 2012; Widome et al., 2009). These changes in dietary intake are commonly attributed to how junk food is a more readily available and calorically dense food that provides rapid satiation suited towards FI populations (Smith et al., 2022). Children experiencing FI may also exhibit a higher likelihood of engaging in binge eating (Bidopia et al., 2023) or emotional eating of highly palatable foods (Hemmingsson, 2018). To investigate these associations in our rodent FI model, we first conducted meal pattern analyses when animals were allowed to freely feed on standard chow shortly after the scheduled feeding period, and again later in adulthood when provided with an array of fatty, sugary, and processed foods designed to mimic a highly palatable junk food diet. While we did not observe any differences in eating behavior or food choice in either of these assessments, Myers and colleagues (2022) found that adult female FI rats have increased total intake and larger first meal sizes when provided with an ad libitum palatable liquid diet.

Estacio and colleagues (2021) also reported increased consumption of a high-fat diet one month after FI had ended in adult female mice. It is thus likely that there are life stage and/or sex-specific effects of FI on feeding behavior.

In addition to altering dietary habits, FI during childhood and adolescence has also been associated with increased mental health disorders as well as emotional, social, and behavioral challenges (Bruening et al., 2017; Burke et al., 2016; Cain et al., 2022; Poole-Di Salvo et al., 2016; Shankar et al., 2017). This includes higher instances of anxiety and hyperactivity, yet there is a lack of literature that tracks these outcomes once adulthood is reached. In the present study, a history of juvenile FI did not produce any differences in anxiety-like behavior during adulthood as assessed in the zero maze and open field tests. The SM group that received HFHS diet on alternating days during scheduled feeding did have increased locomotion compared to the SC group, but IM animals were not significantly different from either group, indicating that FI did not alter locomotor activity. Exposure to high-fat diet has been found to initially induce hyperactivity that leads to hypoactivity after sustained feeding (Wu et al., 2018). It has also been found that limited (2 hours a day, 3 days a week), but not continuous, high fat diet feeding during adolescence produces hyperlocomotion once high fat diet access is stopped, but this effect diminished over time (Blanco-Gandía et al., 2019). It is unclear how unpredictable access to high-fat diet may modulate its ability to induce hyperactivity.

To assess the impact of early life FI on subsequent impulsivity during adulthood, rats were trained to withhold lever presses for increasing time intervals before the press is rewarded with a high-fat sugar-enriched pellet, and no differences were observed between groups over the course of training with regards to either motivation (number of lever presses) or impulsivity (efficiency in reinforcements received relative to lever press number). In human adolescents, FI is associated with attention-deficit hyperactivity disorder symptoms, poor self-control, and future impulse control problems as adults (Jackson et al., 2018; Lu et al., 2019; Vaughn et al., 2016). However, in many cases the assessment of FI status in humans incorporates both inadequate food and unpredictable access to food (and does not distinguish between the two), suggesting that there may be differences between the effects of early life ‘low food security’ (unpredictable food access with adequate food) and ‘very low food security’ (unpredictable food access and inadequate food) that were not captured in our rodent study modeling only low food security.

It is also well documented that early life FI is correlated with developmental delays in cognition, leading to deficits in reading, vocabulary, and mathematic skills and a higher likelihood of repeating a grade (Alaimo et al., 2001; de Oliveira et al., 2020; Gallegos et al., 2021). FI’s association with poorer academic performance is evidenced throughout preschool to teenage years, but there is limited research detailing its long-term impacts on cognition into adulthood. One study surveyed adults aged 50 years or older and found that retrospective self-report of experiencing FI before the age of ten was associated with impaired performance of a word recall memory task (Saenz et al., 2022). In the present study, adulthood cognitive function was assessed in two learning and memory tasks. NOR is a perirhinal cortex-dependent task that tests rodents’ object recognition memory through their ability to discriminate between familiar and novel objects (Albasser et al., 2011). While we observed no group differences in NOR, the IM group showed impaired performance in NLR, which tests hippocampus-dependent spatial memory by evaluating rodents’ exploration preference of an object moved to a new location (Mumby et al., 2002). Unlike the SC and SM control groups, IM (FI-exposed) animals did not demonstrate recognition of the object in a novel location. Myers and colleagues (2022) similarly revealed that NOR performance is not impaired in food-insecure young adult female rats, but a history of FI produced deficits in aging female mice (Estacio et al., 2021). Importantly, there were differences in initial object exploration time, memory retention interval, and testing session length between the NOR assays used in each study, potentially contributing to varying results. NLR has also been investigated in food-insecure juvenile female rats exclusively fed a high-fat diet, but spatial memory impairments were driven by diet composition rather than type of food access (Spaulding et al., 2024). Although NOR and NLR have been separately examined in the context of rodent FI, this is the first paper to test both memory tasks using the same FI model, with NOR selectively engaging perirhinal cortex-dependent memory and NLR engaging hippocampus-dependent memory. Our collective findings provide additional support to previous literature indicating that the hippocampus and associated memory processes are highly vulnerable to dietary insults during early life (Hayes, Kao, et al., 2024). These data also provide evidence that, during juvenile development, unpredictable food access is sufficient to produce memory impairments in adulthood.

One limitation of the present study involved the need to refine the food rations during early stages of the scheduled feeding period. During the first 4-day block of scheduled feeding, nearly every animal in the SC group left some of the day’s rations uneaten, and the SM and IM groups also initially had leftovers on the days they received chow. To mitigate this effect and encourage full consumption of all meals, the 100% baseline food intake trajectory was reduced to 80%. Regardless of this refinement of the model, overall caloric intake did not differ during the scheduled feeding period, confirming that our findings are due to unpredictable food access and not lack of food. A second limitation of this model is the lack of an insecure chow group that would have received the same feeding schedule as the Insecure-Mixed (IM) group except with access to chow only. The inclusion of such a group could help determine whether memory impairments are associated with unpredictable food quantity independent of unpredictable food type.

The FI model developed in this study is the first to our knowledge to simultaneously target three distinct factors of unpredictable food access by manipulating the quantity, timing, and type of food delivered. Our findings reveal that early life food insecurity in the absence of differences in food intake (‘low food security’) impairs hippocampus-dependent spatial recognition memory during adulthood, highlighting the vulnerability of the hippocampus to early life influences. While the absence of a standardized feeding protocol makes it difficult to compare FI’s causal effects between different models, whether during development or adulthood, our novel model allows for improved understanding of the effects of FI with 3 levels of unpredictability, yet absent overall energy restriction. It is important to note that, analogous to the existing rodent models, the experience of FI itself in humans is not universal, and diverse mechanistic animal models can provide insight into how various forms and severities of FI may differentially affect metabolic, cognitive, and behavioral outcomes. Given the apparent sex differences in the impact of FI on metabolic outcomes in both humans and rodent models, we now intend to investigate the impact of our novel model of FI on adult outcomes in females.

## Ethical statement

All experiments reported in this manuscript were approved by the Institutional Animal Care and Use Committee at the University of Southern California (protocol #21096) and performed in accordance with the National Research Council Guide for the Care and Use of Laboratory Animals, which is in compliance with the National Institutes of Health Guide for the Care and use of Laboratory Animals (NIH Publications No. 8023, revised 1978).

## CRediT authorship contribution statement

**Alicia E. Kao**: Conceptualization, Data curation, Formal analysis, Investigation, Methodology, Visualization, Writing – original draft, Writing – review and editing. **Olivia P. Moody**: Data curation, Investigation, Methodology, Writing – review and editing. **Emily E. Noble**: Conceptualization, Writing – review and editing. **Kevin P. Myers**: Conceptualization, Methodology, Resources, Writing – review and editing. **Scott E. Kanoski**: Conceptualization, Funding acquisition, Resources, Supervision, Writing – review & editing. **Anna M. R. Hayes**: Conceptualization, Data curation, Formal analysis, Investigation, Methodology, Visualization, Writing – original draft, Writing – review and editing.

## Data availability

All data associated with this article are available upon reasonable request from the corresponding authors (SEK and AMRH).

## Appendix A. Supplementary information

Supplementary information to this article can be found online.

## Acknowledgements

The authors would like to thank the Kanoski Lab undergraduate research assistants for their assistance with the scheduled feeding protocol and behavioral experiments. Figure subpanels 1A-B, 3A, 3D, 4A, 4D, and 4G were created through Biorender.com.

## References

1. Alaimo, K., Olson, C. M., & Frongillo, E. A., Jr. (2001). Food insufficiency and American school-aged children’s cognitive, academic, and psychosocial development. Pediatrics, 108(1), 44–53.

2. Alamy, M., & Bengelloun, W. A. (2012). Malnutrition and brain development: an analysis of the effects of inadequate diet during different stages of life in rat. Neurosci Biobehav Rev, 36(6), 1463–1480. 10.1016/j.neubiorev.2012.03.009

3. Albasser, M. M., Amin, E., Iordanova, M. D., Brown, M. W., Pearce, J. M., & Aggleton, J. P. (2011). Perirhinal cortex lesions uncover subsidiary systems in the rat for the detection of novel and familiar objects. European Journal of Neuroscience, 34(2), 331–342. 10.1111/j.1460-9568.2011.07755.x

4. Andrews, C., Zuidersma, E., Verhulst, S., Nettle, D., & Bateson, M. (2021). Exposure to food insecurity increases energy storage and reduces somatic maintenance in European starlings (Sturnus vulgaris). R Soc Open Sci, 8(9), 211099. 10.1098/rsos.211099

5. Bateson, M., Andrews, C., Dunn, J., Egger, C., Gray, F., McHugh, M., & Nettle, D. (2021). Food insecurity increases energetic efficiency, not food consumption: an exploratory study in European starlings. PeerJ, 9, e11541. 10.7717/peerj.11541

6. Belzung, C., & Griebel, G. (2001). Measuring normal and pathological anxiety-like behaviour in mice: a review. Behavioural Brain Research, 125(1), 141–149. 10.1016/S0166-4328(01)00291-1

7. Berge, J. M., Fertig, A. R., Trofholz, A., Neumark-Sztainer, D., Rogers, E., & Loth, K. (2020). Associations between parental stress, parent feeding practices, and child eating behaviors within the context of food insecurity. Prev Med Rep, 19, 101146. 10.1016/j.pmedr.2020.101146

8. Bhargava, A., Jolliffe, D., & Howard, L. L. (2008). Socio-economic, behavioural and environmental factors predicted body weights and household food insecurity scores in the Early Childhood Longitudinal Study-Kindergarten. Br J Nutr, 100(2), 438–444. 10.1017/s0007114508894366

9. Bidopia, T., Carbo, A. V., Ross, R. A., & Burke, N. L. (2023). Food insecurity and disordered eating behaviors in children and adolescents: A systematic review. Eat Behav, 49, 101731. 10.1016/j.eatbeh.2023.101731

10. Blanco-Gandía, M. C., Miñarro, J., & Rodríguez-Arias, M. (2019). Behavioral profile of intermittent vs continuous access to a high fat diet during adolescence. Behav Brain Res, 368, 111891. 10.1016/j.bbr.2019.04.005

11. Bruening, M., Dinour, L. M., & Chavez, J. B. R. (2017). Food insecurity and emotional health in the USA: a systematic narrative review of longitudinal research. Public Health Nutr, 20(17), 3200–3208. 10.1017/s1368980017002221

12. Burke, M. P., Martini, L. H., Çayır, E., Hartline-Grafton, H. L., & Meade, R. L. (2016). Severity of Household Food Insecurity Is Positively Associated with Mental Disorders among Children and Adolescents in the United States. J Nutr, 146(10), 2019–2026. 10.3945/jn.116.232298

13. Cain, K. S., Meyer, S. C., Cummer, E., Patel, K. K., Casacchia, N. J., Montez, K., Palakshappa, D., & Brown, C. L. (2022). Association of Food Insecurity with Mental Health Outcomes in Parents and Children. Acad Pediatr, 22(7), 1105–1114. 10.1016/j.acap.2022.04.010

14. Casey, P. H., Simpson, P. M., Gossett, J. M., Bogle, M. L., Champagne, C. M., Connell, C., Harsha, D., McCabe-Sellers, B., Robbins, J. M., Stuff, J. E., & Weber, J. (2006). The association of child and household food insecurity with childhood overweight status. Pediatrics, 118(5), e1406–1413. 10.1542/peds.2006-0097

15. Cohn-Schwartz, E., & Weinstein, G. (2020). Early-life food deprivation and cognitive performance among older Europeans. Maturitas, 141, 26–32. 10.1016/j.maturitas.2020.06.020

16. Dana, L. M., Ramos-García, C., Kerr, D. A., Fry, J. M., Temple, J., & Pollard, C. M. (2025). Social Vulnerability and Child Food Insecurity in Developed Countries: A Systematic Review. Adv Nutr, 16(2), 100365. 10.1016/j.advnut.2025.100365

17. de Oliveira, K. H. D., de Almeida, G. M., Gubert, M. B., Moura, A. S., Spaniol, A. M., Hernandez, D. C., Pérez-Escamilla, R., & Buccini, G. (2020). Household food insecurity and early childhood development: Systematic review and meta-analysis. Matern Child Nutr, 16(3), e12967. 10.1111/mcn.12967

18. Décarie-Spain, L., Hayes, A. M. R., Lauer, L. T., & Kanoski, S. E. (2024). The gut-brain axis and cognitive control: A role for the vagus nerve. Seminars in Cell & Developmental Biology, 156, 201–209. 10.1016/j.semcdb.2023.02.004

19. Denninger, J. K., Smith, B. M., & Kirby, E. D. (2018). Novel Object Recognition and Object Location Behavioral Testing in Mice on a Budget. J Vis Exp(141). 10.3791/58593

20. Dubois, L., Bédard, B., Goulet, D., Prud’homme, D., Tremblay, R. E., & Boivin, M. (2023). Experiencing food insecurity in childhood: influences on eating habits and body weight in young adulthood. Public Health Nutr, 26(11), 2396–2406. 10.1017/s1368980023001854

21. Eisenmann, J. C., Gundersen, C., Lohman, B. J., Garasky, S., & Stewart, S. D. (2011). Is food insecurity related to overweight and obesity in children and adolescents? A summary of studies, 1995-2009. Obes Rev, *12*(5), e73-83. 10.1111/j.1467-89X.2010.00820.x

22. Eskandari, F., Lake, A. A., Rose, K., Butler, M., & O’Malley, C. (2022). A mixed-method systematic review and meta-analysis of the influences of food environments and food insecurity on obesity in high-income countries. Food Sci Nutr, 10(11), 3689–3723. 10.1002/fsn3.2969

23. Estacio, S. M., Thursby, M. M., Simms, N. C., Orozco, V. A., Wu, J. P., Miawotoe, A. A., Worth, W. W., Capeloto, C. B., Yamashita, K., Tewahade, K. R., & Saxton, K. B. (2021). Food insecurity in older female mice affects food consumption, coping behaviors, and memory. PLOS ONE, 16(4), e0250585. 10.1371/journal.pone.0250585

24. Fleming, M. A., Kane, W. J., Meneveau, M. O., Ballantyne, C. C., & Levin, D. E. (2021). Food Insecurity and Obesity in US Adolescents: A Population-Based Analysis. Child Obes, 17(2), 110–115. 10.1089/chi.2020.0158

25. Frongillo, E. A., Adebiyi, V. O., & Boncyk, M. (2024). Meta-review of child and adolescent experiences and consequences of food insecurity. Global Food Security, 41, 100767.

26. Gallegos, D., Eivers, A., Sondergeld, P., & Pattinson, C. (2021). Food Insecurity and Child Development: A State-of-the-Art Review. Int J Environ Res Public Health, 18(17). 10.3390/ijerph18178990

27. Ghadban, E., Fekih-Romdhane, F., Khachan, J., Rizk, M., Ghadbane, C., Mouaness, C., Chehwan, T., El Aam, M., Obeid, S., & Hallit, S. (2025). The relationship between household food insecurity and quality of life among children aged 7-13 years: effects of parent-reported disordered eating, anxiety and depression. BMC Public Health, 25(1), 1008. 10.1186/s12889-025-21785-6

28. Gil, C. R. E., Lund, J., Żylicz, J. J., Ranea-Robles, P., Sørensen, T. I. A., & Clemmensen, C. (2025). Food insecurity promotes adiposity in mice. Obesity (Silver Spring). 10.1002/oby.24259

29. Grutzmacher, S., & Gross, S. (2011). Household food security and fruit and vegetable intake among low-income fourth-graders. J Nutr Educ Behav, 43(6), 455–463. 10.1016/j.jneb.2010.10.004

30. Hayes, A. M. R., Kao, A. E., Ahuja, A., Subramanian, K. S., Klug, M. E., Rea, J. J., Nourbash, A. C., Tsan, L., & Kanoski, S. E. (2024). Early- but not late-adolescent Western diet consumption programs for long-lasting memory impairments in male but not female rats. Appetite, 194, 107150. 10.1016/j.appet.2023.107150

31. Hayes, A. M. R., Lauer, L. T., Kao, A. E., Sun, S., Klug, M. E., Tsan, L., Rea, J. J., Subramanian, K. S., Gu, C., Tanios, N., Ahuja, A., Donohue, K. N., Décarie-Spain, L., Fodor, A. A., & Kanoski, S. E. (2024). Western diet consumption impairs memory function via dysregulated hippocampus acetylcholine signaling. *Brain*, Behavior, and Immunity, 118, 408–422. 10.1016/j.bbi.2024.03.015

32. Hayes, A. M. R., Tsan, L., Kao, A. E., Schwartz, G. M., Décarie-Spain, L., Tierno Lauer, L., Klug, M. E., Schier, L. A., & Kanoski, S. E. (2022). Early Life Low-Calorie Sweetener Consumption Impacts Energy Balance during Adulthood. Nutrients, 14(22), 4709. 10.3390/nu14224709

33. Hemmingsson, E. (2018). Early Childhood Obesity Risk Factors: Socioeconomic Adversity, Family Dysfunction, Offspring Distress, and Junk Food Self-Medication. Curr Obes Rep, 7(2), 204–209. 10.1007/s13679-018-0310-2

34. Hobbs, S., & King, C. (2018). The Unequal Impact of Food Insecurity on Cognitive and Behavioral Outcomes Among 5-Year-Old Urban Children. J Nutr Educ Behav, 50(7), 687–694. 10.1016/j.jneb.2018.04.003

35. Hsu, T. M., Noble, E. E., Liu, C. M., Cortella, A. M., Konanur, V. R., Suarez, A. N., Reiner, D. J., Hahn, J. D., Hayes, M. R., & Kanoski, S. E. (2018). A hippocampus to prefrontal cortex neural pathway inhibits food motivation through glucagon-like peptide-1 signaling. Molecular Psychiatry, 23(7), 1555–1565. 10.1038/mp.2017.91

36. Jackson, D. B., Newsome, J., Vaughn, M. G., & Johnson, K. R. (2018). Considering the role of food insecurity in low self-control and early delinquency. Journal of Criminal Justice, 56, 127–139. 10.1016/j.jcrimjus.2017.07.002

37. Jyoti, D. F., Frongillo, E. A., & Jones, S. J. (2005). Food insecurity affects school children’s academic performance, weight gain, and social skills. J Nutr, 135(12), 2831–2839. 10.1093/jn/135.12.2831

38. Kanoski, S. E., & Grill, H. J. (2017). Hippocampus Contributions to Food Intake Control: Mnemonic, Neuroanatomical, and Endocrine Mechanisms. Biological Psychiatry, 81(9), 748–756. 10.1016/j.biopsych.2015.09.011

39. Lin, W. C., Liu, C., Kosillo, P., Tai, L. H., Galarce, E., Bateup, H. S., Lammel, S., & Wilbrecht, L. (2022). Transient food insecurity during the juvenile-adolescent period affects adult weight, cognitive flexibility, and dopamine neurobiology. Curr Biol, 32(17), 3690–3703.e3695. 10.1016/j.cub.2022.06.089

40. Liu, O. C., Ortiz, R., Baidal, J. W., Pierce, K. A., Perrin, E. M., & Duh-Leong, C. (2025). Childhood Food Insecurity Trajectories and Adult Weight and Self-Reported Health. Am J Prev Med, 107647. 10.1016/j.amepre.2025.107647

41. Lu, S., Perez, L., Leslein, A., & Hatsu, I. (2019). The Relationship between Food Insecurity and Symptoms of Attention-Deficit Hyperactivity Disorder in Children: A Summary of the Literature. Nutrients, 11(3). 10.3390/nu11030659

42. Matheson, D. M., Varady, J., Varady, A., & Killen, J. D. (2002). Household food security and nutritional status of Hispanic children in the fifth grade. Am J Clin Nutr, 76(1), 210–217. 10.1093/ajcn/76.1.210

43. Melchior, M., Chastang, J. F., Falissard, B., Galéra, C., Tremblay, R. E., Côté, S. M., & Boivin, M. (2012). Food insecurity and children’s mental health: a prospective birth cohort study. PLOS ONE, 7(12), e52615. 10.1371/journal.pone.0052615

44. Metallinos-Katsaras, E., Must, A., & Gorman, K. (2012). A longitudinal study of food insecurity on obesity in preschool children. J Acad Nutr Diet, 112(12), 1949–1958. 10.1016/j.jand.2012.08.031

45. Mumby, D. G., Gaskin, S., Glenn, M. J., Schramek, T. E., & Lehmann, H. (2002). Hippocampal damage and exploratory preferences in rats: memory for objects, places, and contexts. Learn Mem, 9(2), 49–57. 10.1101/lm.41302

46. Myers, K. P., Majewski, M., Schaefer, D., & Tierney, A. (2022). Chronic experience with unpredictable food availability promotes food reward, overeating, and weight gain in a novel animal model of food insecurity. Appetite, 176, 106120. 10.1016/j.appet.2022.106120

47. Myers, K. P., & Temple, J. L. (2024). Translational science approaches for food insecurity research. Appetite, 200, 107513. 10.1016/j.appet.2024.107513

48. Nagata, J. M., Chu, J., Cervantez, L., Ganson, K. T., Testa, A., Jackson, D. B., Murray, S. B., & Weiser, S. D. (2023). Food insecurity and binge-eating disorder in early adolescence. Int J Eat Disord, 56(6), 1233–1239. 10.1002/eat.23944

49. Nettle, D., Andrews, C., & Bateson, M. (2017). Food insecurity as a driver of obesity in humans: The insurance hypothesis. Behav Brain Sci, 40, e105. 10.1017/s0140525x16000947

50. Noble, E. E., & Kanoski, S. E. (2016). Early life exposure to obesogenic diets and learning and memory dysfunction. Current Opinion in Behavioral Sciences, 9, 7–14. 10.1016/j.cobeha.2015.11.014

51. Noble, E. E., Wang, Z., Liu, C. M., Davis, E. A., Suarez, A. N., Stein, L. M., Tsan, L., Terrill, S. J., Hsu, T. M., Jung, A. H., Raycraft, L. M., Hahn, J. D., Darvas, M., Cortella, A. M., Schier, L. A., Johnson, A. W., Hayes, M. R., Holschneider, D. P., & Kanoski, S. E. (2019). Hypothalamus-hippocampus circuitry regulates impulsivity via melanin-concentrating hormone. Nat Commun, 10(1), 4923. 10.1038/s41467-019-12895-y

52. Norris, S. A., Frongillo, E. A., Black, M. M., Dong, Y., Fall, C., Lampl, M., Liese, A. D., Naguib, M., Prentice, A., Rochat, T., Stephensen, C. B., Tinago, C. B., Ward, K. A., Wrottesley, S. V., & Patton, G. C. (2022). Nutrition in adolescent growth and development. Lancet, 399(10320), 172–184. 10.1016/s0140-6736(21)01590-7

53. Ortiz-Marrón, H., Ortiz-Pinto, M. A., Urtasun Lanza, M., Cabañas Pujadas, G., Valero Del Pino, V., Belmonte Cortés, S., Gómez Gascón, T., & Ordobás Gavín, M. (2022). Household food insecurity and its association with overweight and obesity in children aged 2 to 14 years. BMC Public Health, 22(1), 1930. 10.1186/s12889-022-14308-0

54. Parent, M. B., Higgs, S., Cheke, L. G., & Kanoski, S. E. (2022). Memory and eating: A bidirectional relationship implicated in obesity. Neurosci Biobehav Rev, 132, 110–129. 10.1016/j.neubiorev.2021.10.051

55. Poole-Di Salvo, E., Silver, E. J., & Stein, R. E. (2016). Household Food Insecurity and Mental Health Problems Among Adolescents: What Do Parents Report? Acad Pediatr, 16(1), 90–96. 10.1016/j.acap.2015.08.005

56. Rabbitt, M. P., Reed-Jones, M., Hales, L. J., & Burke, M. P. (2024). *Household food security in the United States in 2023 (Report No. ERR-337)*. U. S. Department of Agriculture Economic Research Service. 10.32747/2024.8583175.ers

57. Rahi, B., Al Mashharawi, F., Harb, H., El Khoury-Malhame, M., & Mattar, L. (2025). Food Insecurity and Coping Mechanisms: Impact on Maternal Mental Health and Child Malnutrition. Nutrients, 17(2). 10.3390/nu17020330

58. Rea, J. J., Liu, C. M., Hayes, A. M. R., Bashaw, A. G., Schwartz, G. M., Ohan, R., Décarie-Spain, L., Kao, A. E., Klug, M. E., Phung, K. J., Waldow, A. I., Wood, R. I., & Kanoski, S. E. (2025). Hippocampus Oxytocin Signaling Promotes Prosocial Eating in Rats. Biol Psychiatry, 97(5), 540–549. 10.1016/j.biopsych.2024.07.014

59. Reck, A., Sweet, L. H., Geier, C., Kogan, S. M., Cui, Z., & Oshri, A. (2024). Food insecurity and adolescent impulsivity: The mediating role of functional connectivity in the context of family flexibility. Dev Sci, 27(6), e13554. 10.1111/desc.13554

60. Rose, D., & Bodor, J. N. (2006). Household food insecurity and overweight status in young school children: results from the Early Childhood Longitudinal Study. Pediatrics, 117(2), 464–473. 10.1542/peds.2005-0582

61. Saavedra, J. M., & Prentice, A. M. (2023). Nutrition in school-age children: a rationale for revisiting priorities. Nutr Rev, 81(7), 823–843. 10.1093/nutrit/nuac089

62. Saenz, J. L., Kessler, J., & Nelson, E. (2022). Food Insecurity across the Life-Course and Cognitive Function among Older Mexican Adults. Nutrients, 14(7). 10.3390/nu14071462

63. Shankar, P., Chung, R., & Frank, D. A. (2017). Association of Food Insecurity with Children’s Behavioral, Emotional, and Academic Outcomes: A Systematic Review. J Dev Behav Pediatr, 38(2), 135–150. 10.1097/dbp.0000000000000383

64. Sharkey, J. R., Nalty, C., Johnson, C. M., & Dean, W. R. (2012). Children’s very low food security is associated with increased dietary intakes in energy, fat, and added sugar among Mexican-origin children (6-11 y) in Texas border Colonias. BMC Pediatr, 12, 16. 10.1186/1471-2431-12-16

65. Smith, L., Barnett, Y., López-Sánchez, G. F., Shin, J. I., Jacob, L., Butler, L., Cao, C., Yang, L., Schuch, F., Tully, M., & Koyanagi, A. (2022). Food insecurity (hunger) and fast-food consumption among 180 164 adolescents aged 12-15 years from sixty-eight countries. Br J Nutr, 127(3), 470–477. 10.1017/s0007114521001173

66. Spaulding, M. O., Hoffman, J. R., Madu, G. C., Lord, M. N., Iizuka, C. S., Myers, K. P., & Noble,E. E. (2024). Adolescent food insecurity in female rodents and susceptibility to diet-induced obesity. Physiol Behav, 273, 114416. 10.1016/j.physbeh.2023.114416

67. St Pierre, C., Ver Ploeg, M., Dietz, W. H., Pryor, S., Jakazi, C. S., Layman, E., Noymer, D., Coughtrey-Davenport, T., & Sacheck, J. M. (2022). Food Insecurity and Childhood Obesity: A Systematic Review. Pediatrics, 150(1). 10.1542/peds.2021-055571

68. Suarez, A. N., Liu, C. M., Cortella, A. M., Noble, E. E., & Kanoski, S. E. (2020). Ghrelin and Orexin Interact to Increase Meal Size Through a Descending Hippocampus to Hindbrain Signaling Pathway. Biol Psychiatry, 87(11), 1001–1011. 10.1016/j.biopsych.2019.10.012

69. Tagliaferro, A. R., & Levitsky, D. A. (1982). Overcompensation of food intake following brief periods of food restriction. Physiol Behav, 29(4), 747–750. 10.1016/0031-9384(82)90250-5

70. Tsan, L., Décarie-Spain, L., Noble, E. E., & Kanoski, S. E. (2021). Western Diet Consumption During Development: Setting the Stage for Neurocognitive Dysfunction [Review]. Frontiers in Neuroscience, 15. 10.3389/fnins.2021.632312

71. Vaughn, M. G., Salas-Wright, C. P., Naeger, S., Huang, J., & Piquero, A. R. (2016). Childhood Reports of Food Neglect and Impulse Control Problems and Violence in Adulthood. Int J Environ Res Public Health, 13(4), 389. 10.3390/ijerph13040389

72. Widome, R., Neumark-Sztainer, D., Hannan, P. J., Haines, J., & Story, M. (2009). Eating when there is not enough to eat: eating behaviors and perceptions of food among food-insecure youths. Am J Public Health, 99(5), 822–828. 10.2105/ajph.2008.139758

73. Wirth, S. H., Palakshappa, D., & Brown, C. L. (2020). Association of household food insecurity and childhood weight status in a low-income population. Clin Obes, 10(6), e12401. 10.1111/cob.12401

74. Wu, H., Liu, Q., Kalavagunta, P. K., Huang, Q., Lv, W., An, X., Chen, H., Wang, T., Heriniaina, R. M., Qiao, T., & Shang, J. (2018). Normal diet Vs High fat diet - A comparative study: Behavioral and neuroimmunological changes in adolescent male mice. Metab Brain Dis, 33(1), 177–190. 10.1007/s11011-017-0140-z

75. Yang, M., Singh, A., de Araujo, A., McDougle, M., Ellis, H., Décarie-Spain, L., Kanoski, S. E., & de Lartigue, G. (2025). Separate orexigenic hippocampal ensembles shape dietary choice by enhancing contextual memory and motivation. Nat Metab, 7(2), 276–296. 10.1038/s42255-024-01194-6

76. Zhong, D., Gunnar, M. R., Kelly, A. S., French, S., Sherwood, N. E., Berge, J. M., & Kunin- Batson, A. (2022). Household food insecurity and obesity risk in preschool-aged children: A three-year prospective study. Soc Sci Med, 307, 115176. 10.1016/j.socscimed.2022.115176

